# The Subjective Value of Cognitive Effort is Encoded by a Domain-General Valuation Network

**DOI:** 10.1101/391805

**Authors:** Andrew Westbrook, Bidhan Lamichhane, Todd Braver

## Abstract

Cognitive control is necessary for goal-directed behavior, yet people treat control as costly, discounting goal value by cognitive demands in a similar manner as they would for delayed or risky outcomes. It is unclear, however, whether a putatively domain-general valuation network implicated in other cost domains also encodes the subjective value (SV) of cognitive effort. Here, we demonstrate that a valuation network, centered on the ventromedial prefrontal cortex and ventral striatum, also encodes SV during cognitive effort-based decision-making. We doubly dissociate this network from a primarily frontoparietal network recruited as a function of decision difficulty. We also find evidence that SV signals predict choice and are influenced by state and trait motivation, including sensitivity to reward and anticipated task performance. These findings unify cognitive effort with other cost domains, and inform physiological mechanisms of SV representations underlying the willingness to expend cognitive effort.

## Introduction

Cognitive control is required for flexible, precise, goal-directed behavior (Botvinick et al., 2001; Egner and Hirsch, 2005; Miller, 2000). Yet, intriguingly, control is subjectively costly, leading individuals to avoid control demands, even when foregoing valuable outcomes (Dixon and Christoff, 2012; Kool et al., 2010; Westbrook et al., 2013). Subjectively high effort costs have clinical consequences, undermining functioning in disorders as diverse as schizophrenia (Culbreth et al., 2016; Gold et al., 2015), ADHD (Volkow et al., 2010), depression (Cohen et al., 2001), and Parkinson’s disease (Manohar et al., 2015; Sinha et al., 2013). It is thus critical to identify cognitive effort-based decision-making mechanisms, to address motivational impairments in cognitive function and develop new targets for clinical intervention.

Effort-based decision-making may involve computation of subjective value (SV) in a “common currency”, by integrating effort costs with reward benefits (Padoa-Schioppa, 2011). Such SV representations would facilitate fungible exchange across cost and benefit dimensions (Rangel et al., 2008). Numerous studies have identified a core valuation network that appears to encode SV, including the ventromedial prefrontal cortex (vmPFC) and ventral striatum (VS) (Bartra et al., 2013; Levy and Glimcher, 2012). This network has been implicated in integrating diverse benefits, including both primary and secondary rewards, with diverse cost factors, such as risk, delay, and physical effort. As yet, however, this network has not been shown to encode SV for choices about cognitive effort.

Indeed, only two prior studies have investigated SV encoding during cognitive effort decisionmaking (Chong et al., 2017; Massar et al., 2015). Surprisingly, they did not implicate the core valuation network. Instead, they identified frontoparietal regions more typically associated with cognitive control itself: the anterior cingulate cortex (ACC), dorsolateral prefrontal cortex (dlPFC), and intra-parietal sulcus (IPS). The involvement of such regions is consistent with an account by which greater working memory and cognitive control resources are recruited to support more difficult decisions: when options are close in value (Pochon et al., 2008; Shenhav et al., 2014). It is also consistent with the specific comparator hypothesis postulated for regions including the ACC (Hunt et al., 2012; Klein-Flugge et al., 2016), in which the differences in chosen versus unchosen offer costs and benefits are tracked. Thus, these prior studies provide valuable novel evidence implicating a dorsal frontoparietal network in decision-making about cognitive effort, while also leaving open the question of whether the SV of cognitive effort is encoded by a ventral, core valuation network.

The fact that these prior studies found no evidence to support SV encoding in the core valuation network raises the possibility cognitive effort-based decision-making involves different mechanisms than other decision-making domains. Perhaps, for example, decisions about cognitive effort engagement are not made by computing SV *per se*, but by prospectively simulating the relative cognitive demands of tasks under consideration, and potential benefits, to determine whether to allocate limited resources. This approach would be consistent with the hypotheses that: 1) the subjective cognitive effort reflects context-specific opportunity costs, proportional to the resources demanded by a given task (Kurzban et al., 2013; Shenhav et al., 2017), and 2) cognitive control is recruited in proportion to the controllability of a task and the expected value of its allocation (Boureau et al., 2015; Musslick et al., 2015; Shenhav et al., 2013).

Alternatively, it may be that the prior cognitive effort studies used methods which were primarily sensitive to detecting regions as a function of choice difficulty rather than single offer SV. Indeed, the primary regressor of interest in one study (Chong et al., 2017) was the difference in offer values, while the value of the discounted offer, orthogonalized with respect to the undiscounted offer was the primary regressor in the other (Massar et al., 2015). Hence, it is critical to complement these approaches with methods focused on isolating the representation of SV of individual offers. Furthermore, to demonstrate SV encoding, it is critical to show that SV representations are simultaneously sensitive to all choice features including both costs (cognitive load) and benefits (reward magnitude). Regions tracking either of these dimensions alone would correlate with SV, but not encode SV *per se*. Reward processing is ubiquitous in the brain – diverse regions are increasingly active in response to cues of increasing reward magnitude (Vickery et al., 2011). Similarly, regions like the dorsal ACC (dACC) are well-established to encode anticipated cognitive demand (Kerns, 2004; Ridderinkhof et al., 2004), and may thus correlate with, but not strictly encode SV. Alternatively, given its hypothesized role in computing the expected value of control, the dACC may track both costs and benefits (Shenhav et al., 2013). Finally, it is noteworthy that although many neuroeconomic studies show BOLD signal tracking objective cost-benefit features, few studies test whether putative SV representations are truly subjective. Claiming that a region tracks *subjective* value, requires demonstrating that value signals covary with subjective measures of objective choice features. Note that subjectivity may reflect either stable, trait differences, context-specific state differences, or both (Westbrook and Braver, 2015). For example, state willingness to exert cognitive effort is known to vary as a function of sleep deprivation (Libedinsky et al., 2013).

In the present study, we utilized a variant of a cognitive effort discounting task (COGED) (Westbrook and Braver, 2015), in conjunction with fMRI, to test whether a domain-general valuation network, by comparison with regions associated with cognitive control implicated in two prior studies, encodes the SV of cognitive effort. In prior work, we have shown that COGED choices are sensitive to both objective features like cognitive load and reward, and subjective, trait features like typical daily engagement with cognitive demands (Westbrook et al., 2013), cognitive aging (Westbrook et al., 2013), and negative symptoms in schizophrenia (Culbreth et al., 2016). Here, we modified COGED to test whether a putatively domain-general valuation network encodes SV during evaluation of single offers, thus isolated from decision-making processes and decision difficulty effects. We further tested whether regions encoding SV reflected all three cardinal dimensions: costs, benefits, and subjectivity. Finally, we examined factors giving rise to subjectivity, as well as the relationship between SV representations and ultimate choice outcomes.

## Results

### Reward is Discounted Subjectively by Cognitive Load

In COGED, participants decide whether to perform variously demanding N-back working memory task levels for money (Braver et al., 1997; Kirchner, 1958). Prior work has demonstrated that the N-back is perceived as effortful, and self-reported effort increases systematically with cognitive load (Hopstaken et al., 2015; Westbrook et al., 2013). Critically, subjective effort costs are measured by the extent to which cash offers are discounted for each load level (N) (Westbrook et al., 2013). In the present study, outside the scanner, participants first experienced all N-back levels, then made repeated decisions between performing a high-load N-back level (N = 2–6) for one of three base offer amounts ($2, $3, or $4) or instead performing the low-load 1-back condition for a smaller, variable amount. Low-load (1-back) offers were iterated in a staircase fashion, until an indifference point was reached. Indifference points indicate the subjective value (SV) of offers, discounted by costs of each higher N-back load relative to the low-load baseline. Replicating prior findings (Westbrook et al., 2013), offers were discounted at all (high) load levels (SV < 1), with SV reliably decreasing as load increased (F_1,20_ = 49.3; p < 0.01; η^2^ = 0.68), indicating rising effort costs (Figure 1). Importantly, there were strong individual differences in the degree of discounting. Given theoretical uncertainty regarding the form of the discount function (Chong et al., 2017; Hartmann et al., 2013), we quantified individual differences with an area under the curve (AUC; Figure 1) measure that prior work has shown to be psychometrically optimal for individual difference analyses (Myerson et al., 2001). Higher AUC indicates that a participant is more willing to expend cognitive effort for reward, on average.

**Figure 1.**
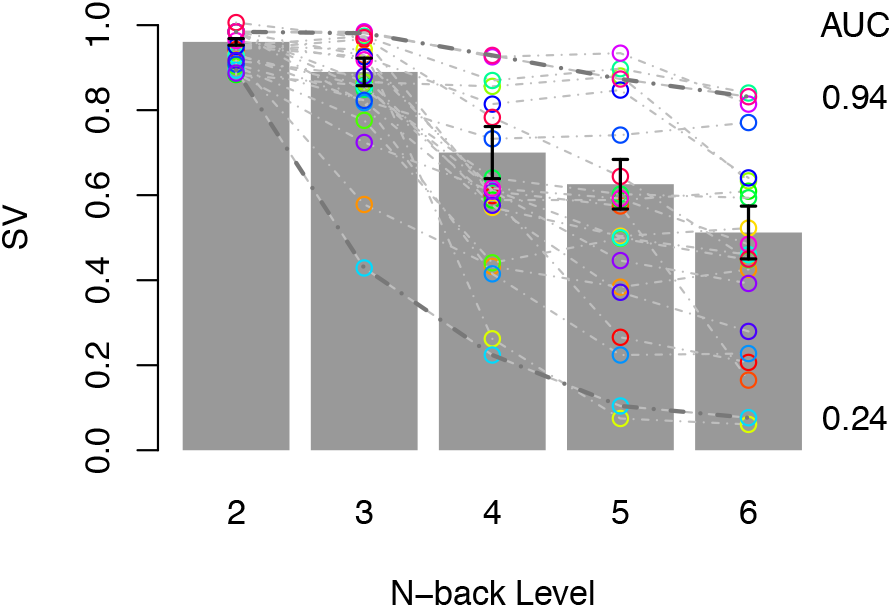
Decreasing SV reflects rising subjective costs as N-back level increases. Grey bars and black lines reflect group means and SEM. Grey dashed lines show individual participants’ discounting curves. Area Under the Curve (AUC) values provided for two example participants.

Participants may discount high-effort tasks because they anticipate poorer performance as task load increases. However, it is unlikely that discounting reflects performance alone. First, participants were instructed that they would be paid for completing a task, even if they performed poorly. Second, although poor performance predicts steeper discounting, there is considerable discounting variance not explained by performance. For example, even when controlling for N-back performance (d′; B = 9.83×10^-2^, p = 0.02), load significantly (B = −8.16×10^-2^, p < 0.01) predicted SV in a hierarchical multiple regression (load levels nested within participants). Furthermore, indexed by d′, steep discounters (below-median AUC) performed the N-back as well as shallow discounters (Wilcoxon p = 0.62), and numerically better at high load levels (though not reliably: p’s ≥ 0.16). Hence, although declining performance with higher load may contribute to discounting, it does not satisfactorily explain individual differences.

In the fMRI scanner, participants again decided between a base offer ($2, $3, or $4) to perform the high-load N-back (N = 2–6), and a variable amount for the low-load, but this time with the low-load offer systematically adjusted with respect to participants’ own indifference points, to control decision difficulty and balance choice bias. Specifically, with *γ* referring to the fractional difference between the indifference offer and the bounds of $0 or the base amounts ($2, $3, or $4), participants decided between offers in which the low-load amount was slightly above indifference (*γ* = 0.2, 0.6), biasing low-load choices (“low-load biased”), or slightly below indifference (*γ* = −0.1, −0.4), biasing high-load choices (“high-load biased” trials). We also included *catch* trials, in which we offered equal amounts (*γ* = 1.0) for the 1-back and high-load task, strongly biasing low-load choices (“low-load catch”) or instead $0 (*γ* = −1.0), strongly biasing high-load choices (“high-load catch”). A key advantage of this design is that decision difficulty, high-load versus low-load preference, and SV were all orthogonalized across trials.

As anticipated, bias reliably (F_1,20_ = 341.1; p < 0.01) influenced the probability of selecting the high-load offer on a given trial. Moreover, even excluding catch trials, there was a reliable preference for high-load offers on high-load biased trials (choice probability > 0.5; t_20_ = 5.12; p < 0.01), and a reliable preference for low-load offers on low-load biased trials (*γ* > 0; choice probability < 0.5; t_20_ = 2.31; p = 0.03; Figure 2A).

**Figure 2.**
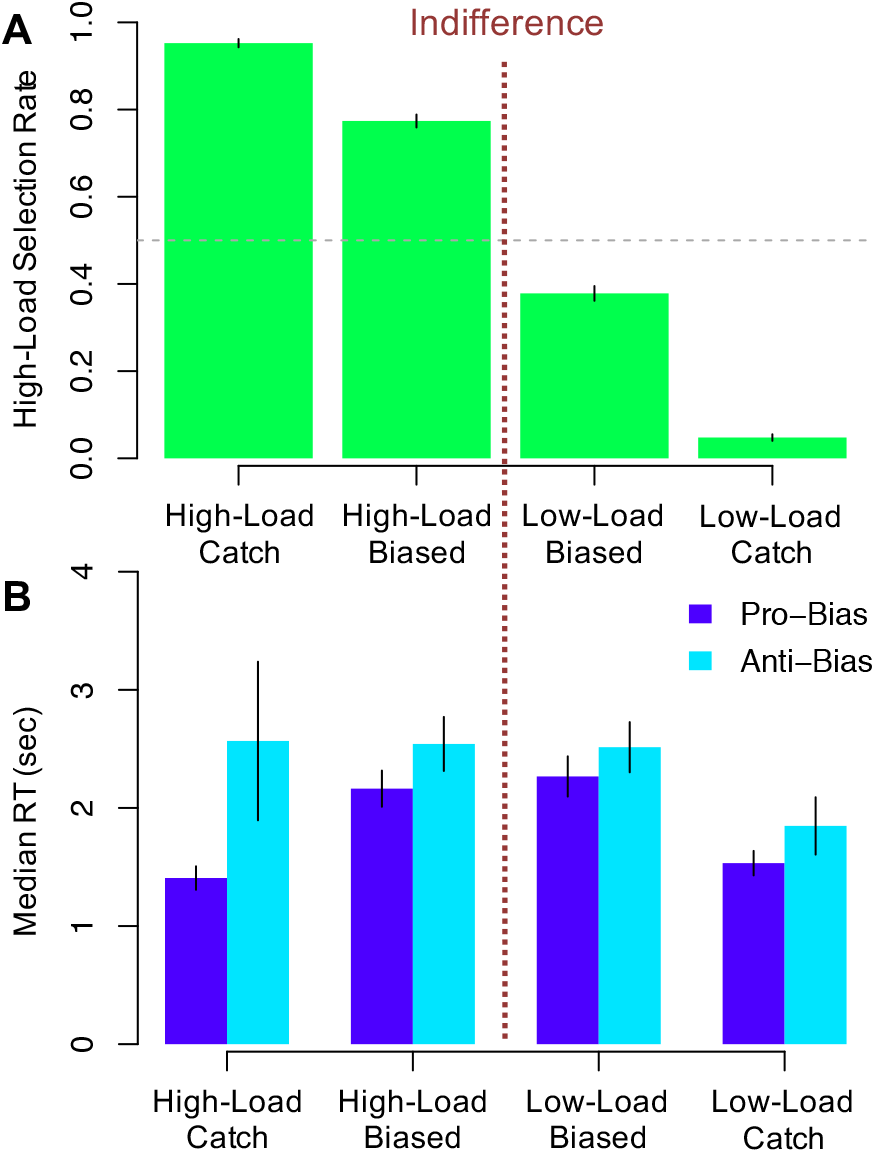
A) Proportion of high demand options selected as a function of biasing of the low demand offer, and N-back level. B) Median reaction times as a function of proximity and whether choices went with (pro-) or against (anti-) low demand offer biasing (High-Load: *γ* < 0; Low-Load: *γ* > 0).

Choice reaction times provide further evidence that our discounting procedure identified indifference points accurately: participants responded more slowly on trials closer to indifference, consistent with increasing decision difficulty. On “pro-bias” trials, in which participants’ choices were consistent with offer biases, median reaction times were slower on more difficult decision trials (|*γ*| < 1.0) relative to catch trials (*γ* = −1.0 or 1.0; t_20_ = 7.51, p < 0.01; Figure 2B). Furthermore, when decisions went against offer biases (“anti-bias” trials), reaction times were significantly slower than on pro-bias trials (t_20_ = 7.03, p < 0.01). Such a pattern is consistent with a response conflict account of decision difficulty (Botvinick, 2007; Pochon et al., 2008; Yarkoni et al., 2005), as conflict would be highest, on average, when deciding against typically preferred alternatives.

### SV Encoding

Our central question was whether a domain-general valuation network encoded cognitive effort-discounted SV. To test this, we asked whether BOLD response reliably tracked the SV of offers to repeat N-back tasks for money. Specifically, participants were instructed to consider the value of single high-load N-back options (N= 2—6) paired with single base amounts ($2, $3, or $4), presented in isolation for 6 seconds (i.e. high-load offers were presented first; Figure 3). Consequently, brain activity 6—8 s after first offer onset was regarded as reflecting single offer valuation, temporally isolated from other decision processes, accounting for hemodynamic lag (Miezin et al., 2000). We tested whether activity during this “valuation period” tracked trial-by-trial, first-offer SV within a set of regions of interest (ROIs) derived from prior meta-analyses identifying the putatively domain-general valuation network (yellow in Figure 4A) including the vmPFC, VS, anterior insula (AI), posterior cingulate cortex (PCC), and dACC (Bartra et al., 2013; Levy and Glimcher, 2012). The focus on *a priori* ROIs was motivated by strong prior beliefs about regions encoding SV, and the desire to maximize statistical power (for completeness, a whole-brain analysis is provided in Supplementary Figure S.1, although this did not identify any regions outside of our candidate ROIs). We contrasted the encoding of SV in this domain-general network with sets of candidate loci identified in two recent studies specifically focused on decision-making about cognitive effort (blue in Figure 4A) (Chong et al., 2017; Massar et al., 2015). Note that to combine ROIs as a set in an unbiased way (each ROI gets equal weight), the response of each ROI at 6—8s was z-scored across trials and then ROIs were averaged together to obtain a network-level response on each trial.

**Figure 3.**
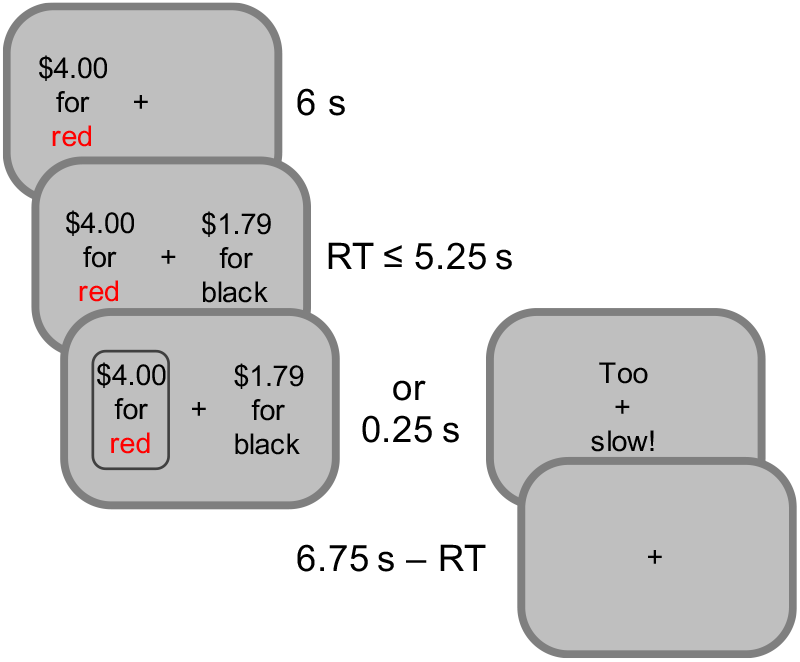
Imaging trials begin with a high-amount, high-load offer (load is indicated by a color label, e.g. red for 2-back). After 6s the 1-back (black task) offer is presented. Participants have 5.25s to respond. After response is indicated briefly, a fixation cross is presented until the end of the trial.

**Figure 4.**
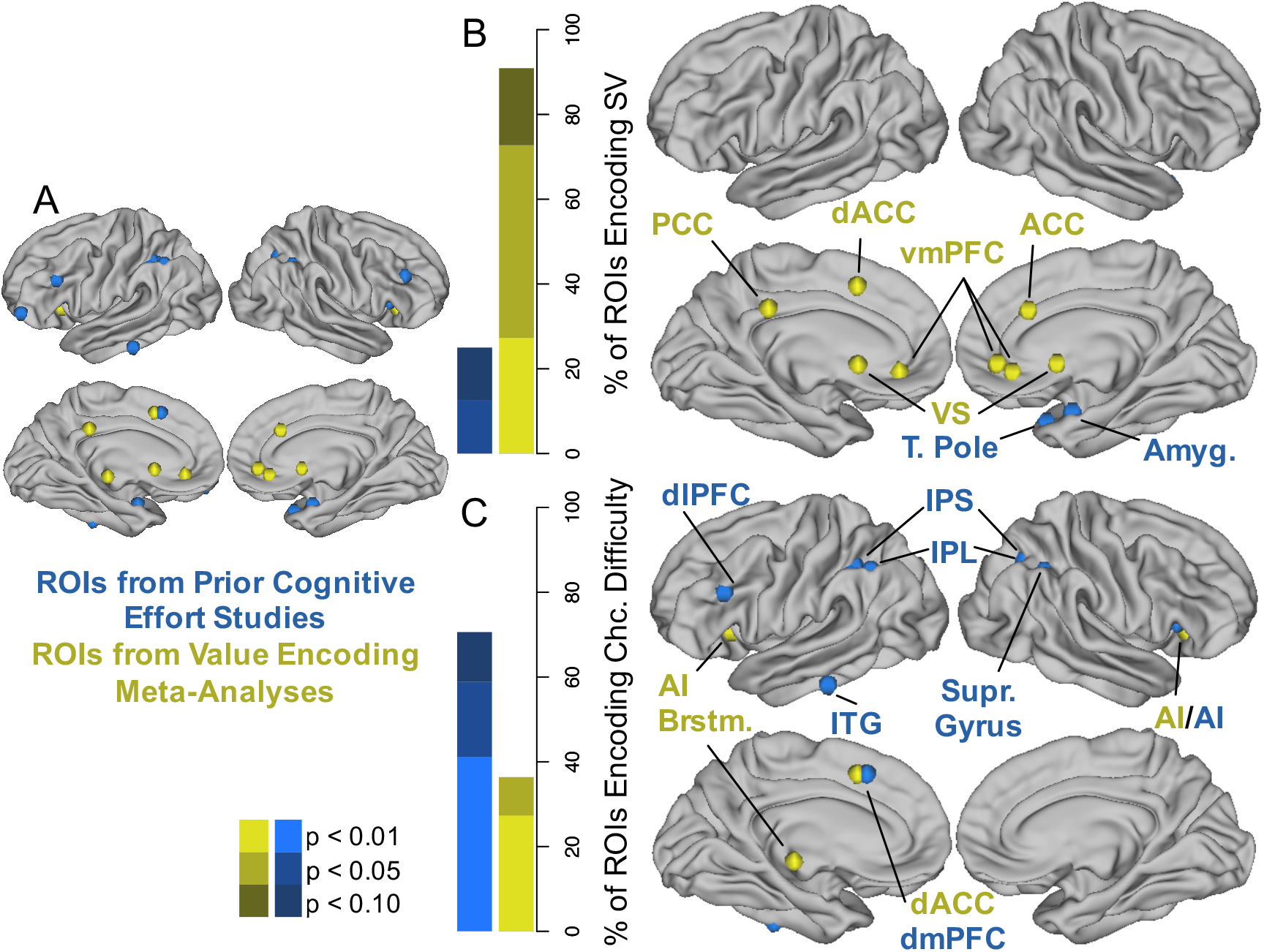
A. 6 mm radius spheres centered at all a *priori* ROIs. Colors indicate origination from either of two prior meta-analyses of domain-general SV encoding (yellow^15,16^) or two recent studies on SV encoding during cognitive effort decision-making (blue^17,18^). Note that the amygdala is projected to the surface for display purposes only. B. Proportion of ROIs from each set tracking SV 6—8s after first offer onset by p-value. Map shows ROIs reliably tracking SV at p < 0.05. C. Proportion of ROIs from each set with reliably more activity on difficult, regular versus easy catch trials by p-value. Map shows ROIs with a reliable difficulty contrast at p < 0.05.

As predicted, the SV meta-analysis network of regions positively and reliably tracked first-offer SV in the valuation period, 6—8 sec following the first offer onset (B = 4.19×10^−2^; p = 0.002). Individual ROIs tracking first offer SV (at p < 0.05) included the bilateral vmPFC, VS, PCC, ACC, and dACC/pre-SMA (Figure 4B; Table 1, Column 3). These results are consistent with the hypothesis that a core valuation network encodes SV, discounted by cognitive effort costs. Furthermore, they extend the notion of domain-generality from delay, risk, and physical effort costs (Bartra et al., 2013; Levy and Glimcher, 2012) to the domain of cognitive effort. At the level of individual ROIs, only the brainstem showed no evidence of encoding SV, despite this region reliably encoding SV in other cost domains (Bartra et al., 2013).

**Table 1.** Effects of choice difficulty and first offer SV, amount and load on BOLD signal in all *a priori* ROIs, grouped by study of origin. Column 2: MNI coordinates for each ROI. Columns 3—5: Effect estimates and corresponding p-values for activity in *a priori* networks and individual ROIs predicting mean residual activity 6—8 s after first offer onset, after regressing out motion and other predictors of non-interest. Column 3 describes first offer SV as a predictor of residuals. Columns 4 and 5 provide relationships with first offer amount and load simultaneously estimated in a hierarchical multiple regression. Column 6: t-tests contrasting the canonical hemodynamic response on regular versus catch trials. Cells are shaded in greyscale according to their p-value. Dark grey is p < 0.01; medium is 0.01 < p < 0.05; light is 0.05 < p < 0.10. Network-level effects at the top of each section were estimated from the mean response at 6—8 s, z scored across trials, then averaged across ROIs in each network.

| Anatomical Description | MNI (x,y,z) | SV | Amount | Load | Regular vs. Catch $t_{20}(p)$ |
| --- | --- | --- | --- | --- | --- |
| | | $B \times 10^{-2}$ (p-value) | | | |
| All SV Meta-Analysis ROIs |  | 4.2 (<0.01) | 4.1 (<0.01) | -5.0 (<0.01) | 1.77 (0.09) |
| Levy and Glimcher (2012) |  |  |  |  |  |
| l vmPFC | (-7,38,-11) | 2.2 (<0.01) | 2.1 (<0.01) | -2.1 (<0.01) | -1.38 (0.18) |
| r vmPFC | (4,35,-12) | 2.2 (<0.01) | 1.8 (<0.01) | -2.3 (<0.01) | -1.22 (0.24) |
| Bartra, McGuire, and Kable (2013) |  |  |  |  |  |
| PCC | (-2,-35,31) | 1.9 (0.04) | 1.7 (0.04) | -2.2 (<0.01) | 0.36 (0.72) |
| dACC/pre-SMA | (-1,15,44) | 2.3 (0.02) | 0.9 (0.28) | -2.5 (<0.01) | 5.21 (<0.01) |
| ACC | (0,30,27) | 1.9 (0.04) | 1.7 (0.04) | -2.2 (<0.01) | 1.19 (0.25) |
| l striatum | (-12,12,-6) | 3.8 (0.02) | 2.5 (<0.01) | -1.8 (<0.01) | 0.92 (0.37) |
| r striatum | (12,10,-6) | 4.1 (0.05) | 2.8 (<0.01) | -1.8 (0.02) | 0.39 (0.70) |
| r vmPFC | (2,46,-8) | 3.1 (<0.01) | 2.4 (0.01) | -3.2 (<0.01) | -1.48 (0.16) |
| l AI | (-30,22,-6) | 0.8 (0.10) | 0.5 (0.26) | -1.2 (<0.01) | 3.28 (<0.01) |
| r AI | (32,30,-6) | 0.8 (0.05) | 0.5 (0.18) | -0.8 (0.02) | 2.28 (0.03) |
| Brainstem | (-2,-22,-12) | 0.9 (0.30) | 1.1 (0.19) | -1.6 (0.05) | 2.97 (<0.01) |
| Massar, Libedinsky, Weiyan, Huettel, and Chee (2015) |  |  |  |  |  |
| All Massar et al. ROIs (Study1) |  | 2.6 (0.05) | 1.8 (0.19) | -3.7 (0.02) | 3.50 (<0.01) |
| r supramarg. gyr. | (33,-52,32) | 0.4 (0.38) | 0.1 (0.81) | -0.7 (0.09) | 2.83 (0.01) |
| l cingulate | (-24,-49,36) | 0.4 (0.24) | 0.1 (0.63) | -0.5 (0.13) | 2.61 (0.02) |
| l inf. temp. gyr. | (-58,-35,-22) | 1.1 (0.08) | 0.5 (0.25) | -1.7 (<0.01) | -1.78 (0.09) |
| l IFG | (-43,53,-4) | 0.4 (0.61) | 0.4 (0.67) | -1.8 (0.03) | 0.66 (0.51) |
| l IPL | (-30,-43,43) | 0.5 (0.35) | 0.3 (0.48) | -0.7 (0.12) | 5.10 (<0.01) |
| l IPL | (-41,-55,46) | -0.2 (0.82) | 0.1 (0.93) | -0.9 (0.22) | 3.42 (<0.01) |
| r temporal pole | (34,16,-26) | 3.1 (0.03) | 1.8 (0.02) | -1.4 (0.08) | 0.04 (0.97) |
| Chong, Apps, Giehl, Sillence, Grima, Husain (2017) |  |  |  |  |  |
| All Chong et al. ROIs (Study2) |  | 2.1 (0.14) | 1.7 (0.20) | -3.5 (0.04) | 3.41 (<0.01) |
| l amygdala | (-24,0,-22) | 2.2 (0.08) | 1.6 (0.04) | -1.7 (0.04) | -0.41 (0.69) |
| r amygdala | (24,0,-22) | 2.8 (0.03) | 2.1 (0.02) | -1.4 (0.13) | -0.15 (0.88) |
| l IPS | (-36,-44,38) | 0.4 (0.50) | 0.1 (0.90) | -0.8 (0.12) | 6.04 (<0.01) |
| dACC/dmPFC | (-4,22,44) | 1.1 (0.12) | 0.4 (0.56) | -1.9 (<0.01) | 2.61 (0.02) |
| r IPS | (24,-60,42) | 0.6 (0.25) | 0.3 (0.56) | -1.0 (0.04) | 4.95 (<0.01) |
| r Insula | (34,22,2) | 0.2 (0.75) | 0.2 (0.65) | -0.4 (0.30) | 3.78 (<0.01) |
| l dIPFC | (-44,26,26) | 0.9 (0.23) | 0.8 (0.18) | -0.6 (0.36) | 3.29 (<0.01) |
| r dIPFC | (44,36,30) | 0.1 (0.95) | -0.4 (0.55) | -1.3 (0.07) | 1.68 (0.11) |
| l Insula | (-28,22,4) | 0.5 (0.17) | 0.3 (0.24) | -0.5 (0.11) | 3.08 (0.01) |

By contrast, sets of ROIs identified by the two prior cognitive effort studies did not reliably encode first-offer SV. The set of ROIs from one study (Study1) encoded SV at trend-level (B = 2.54×10^−2^; p = 0.053) while the set identified in by the other study did not (Study2; B = 2.12×10^−2^; p = 0.14). Moreover, at the sub-network level, most individual ROIs from either study did not reliably track first-offer SV. Notable exceptions to this pattern include the right temporal pole from Study1 and right amygdala from Study2 (the left amygdala also encoded trial wise SV at trend-level). The amygdala is notable as a region that has been previously implicated in encoding SV during cognitive, relative to physical effort-based decision-making (Chong et al., 2017) and also supporting cognitive effort-based decision-making in rats (Hosking et al., 2014). Despite these exceptions, the broader pattern did not replicate the findings from the two prior cognitive effort studies: in neither the intraparietal sulcus (IPS), inferior parietal lobule (IPL), nor the lateral PFC loci did activity reliably track first offer SV.

Direct comparisons between sets of ROIs reveal that the meta-analysis network not only tracks trial-wise SV more reliably than ROIs identified in the two prior cognitive effort studies, but their activity is modulated more strongly by, and in turn explains more trial-by-trial variance in, first offer SV. First, as a group, meta-analysis regression weights (Table 1, Column 3) are reliably larger across individual ROIs than those from the two prior cognitive effort studies (t_20_ = 2.94; p < 0.01). Second, in hierarchical, nested model comparisons, meta-analysis ROIs explained more variance in first-offer SV when added to the full set of ROIs from the prior effort studies. For example, after controlling for activity 6—8 s after first offer onset in all ROIs from Study1 and Study2, adding activity from the left 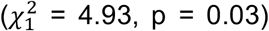 or right 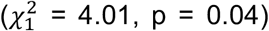 vmPFC, or left 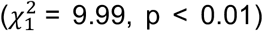 or right 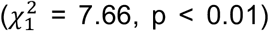 VS significantly improved model deviance. When adding all 11 meta-analysis ROIs at once, model deviance also improved at trend-level 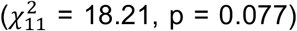 above a model including all ROIs from Study1 and Study2. By contrast, adding Study1 and Study2 ROIs, to a base model containing metaanalysis ROIs, did not improve model deviance 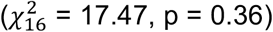.

Relatively weak SV encoding among ROIs from the prior cognitive effort studies suggests that they were previously implicated because they are primarily responsive to factors correlated with SV, rather than SV *per se*. One possibility is decision difficulty. Correlation between SV and difficulty might occur, for example, if participants typically prefer high-effort, high-reward alternatives (such alternatives have higher SV on most trials), and the decision only becomes difficult on infrequent trials when the SV of the high-effort option is low and differences in SV between offers are small. Moreover, as noted, the prior cognitive effort studies also used methods that were likely more sensitive to choice difficulty (e.g., using value difference regressors, or examining BOLD signal while participants contrast two offers rather than evaluate a single offer), rather than single-offer SV.

To examine whether ROIs from the prior cognitive effort studies were relatively more sensitive to decision difficulty, we first tested whether they were more sensitive to differences in offer SV. Specifically, we contrasted hemodynamic response functions, time-locked to second offer onset, between difficult trials, when the differences in SV of were small (|*γ*| < 0.6), and easy catch trials, when differences in SV were large (*γ* = −1.0 or 1.0). In a reversal of the SV encoding analysis, sets of ROIs from the prior cognitive effort studies were robustly sensitive to this difficulty contrast (Study1: t_20_ = 3.50, p < 0.01; Study2: t_20_ = 3.41, p < 0.01) while the SV meta-analysis network was only sensitive at trend-level (t_20_ = 1.77, p = 0.09). The pattern of results across sets of ROIs suggests a double dissociation, which is buttressed by a striking pattern at the individual ROI level: individual ROIs were either more reliably active on difficulty trials or reliably tracked SV, but not both (Table 1; Figure 4B&C). Sole exceptions to this pattern were the dACC and trend-level results in the inferior temporal gyrus and bilateral AI. As noted, ROIs sensitive to the difficulty contrast comprise regions more typically associated with cognitive control, working memory, and evidence accumulation (Basten et al., 2010; Braver et al., 1997; Dosenbach et al., 2006; Kouneiher et al., 2009), including the bilateral IPS, the dACC and pre-supplementary motor area (pre-SMA), and dlPFC. These ROI-level analyses are consistent with a double dissociation, in which a domain-general valuation network tracks first-offer SV significantly better than frontoparietal regions implicated in prior cognitive effort studies, while conversely these latter frontoparietal regions are more sensitive to decision difficulty than those involved with domain-general valuation. As above, for completeness, we examined the decision difficulty contrast across the whole brain; the results (Supplemental Figure S.2; Table S.1) re-capitulate our ROI analysis: a network of regions including the dACC, dlPFC, and IPS were more active on difficult versus easy trials while valuation network regions were not differentially active across trial types.

RT—BOLD signal relationships further support the hypothesis that the prior cognitive effort studies implicated regions that were primarily sensitive to decision difficulty. Namely, mirroring the hierarchical multiple regression strategy above, we tested whether mean BOLD signal 6—8 s after second offer onset (during the decision period) among ROIs from the prior cognitive effort studies explained more variance in RT on pro-bias trials (related to response slowing, when participants’ choices were consistent with their prior discounting patterns, cf. Figure 2B). Adding all 16 ROIs from Study1 and Study2, as a group, improved model deviance 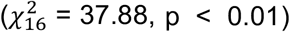 over a base model with only SV meta-analysis ROIs in predicting inverse RT. Conversely, adding the SV meta-analysis ROIs second also improved model deviance 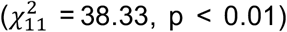, but this is perhaps unsurprising given that these ROIs included the dACC, which is known to robustly track RT across diverse perceptual and economic decision-making tasks (Yarkoni et al., 2009). Indeed, after excluding the dACC, we find that adding SV metaanalysis ROIs second no longer improved model deviance 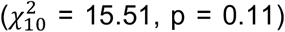, supporting the hypothesis that ROIs identified by the two prior cognitive effort studies are relatively more sensitive to decision difficulty, as indexed by response slowing.

### Costs and Benefits are Jointly Encoded in the Valuation Network

For a region to track SV, it should encode both offer benefits and costs, and with opposing signs. Surprisingly few SV encoding studies decompose these sources of variance, however, leaving open the possibility that BOLD signal correlating with SV may track costs or benefits alone. Thus, we tested whether trial wise first-offer amount (benefits) and N-back load (costs) jointly and independently predicted mean valuation period (at 6—8 s) activity in *a priori* ROIs, again using hierarchical linear models. We found that the SV meta-analysis network covaried both positively with amount (controlling for load; B = 4.07×10^−2^; p = 0.002), and negatively with load (controlling for amount; B = −4.95×10^−2^; p < 0.001; Table 1). This was also true, moreover, among most individual ROIs within this network. For example, bilateral vmPFC and VS activity both reliably increased with higher offer amounts and decreased with increasing task loads (Figure 5). This result supports that the putatively domain-general valuation network not only correlates with SV, but independently tracks both cognitive costs and benefits. By contrast, and mirroring our SV analysis, we did not find encoding of both dimensions in the networks of regions implicated by the prior cognitive effort studies. In both cases, we found evidence that these networks encoded first-offer load (Study1: B = −3.70×10^−2^, p = 0.02; Study2: B = −3.51×10^−2^, p = 0.04), but not first-offer amount (Study1: B = 1.77×10^−2^, p = 0.19; Study2: B = 1.67×10^−2^, p = 0.20). Relative insensitivity to first offer amount helps explain why these ROIs did not reliably track first-offer SV.

**Figure 5.**
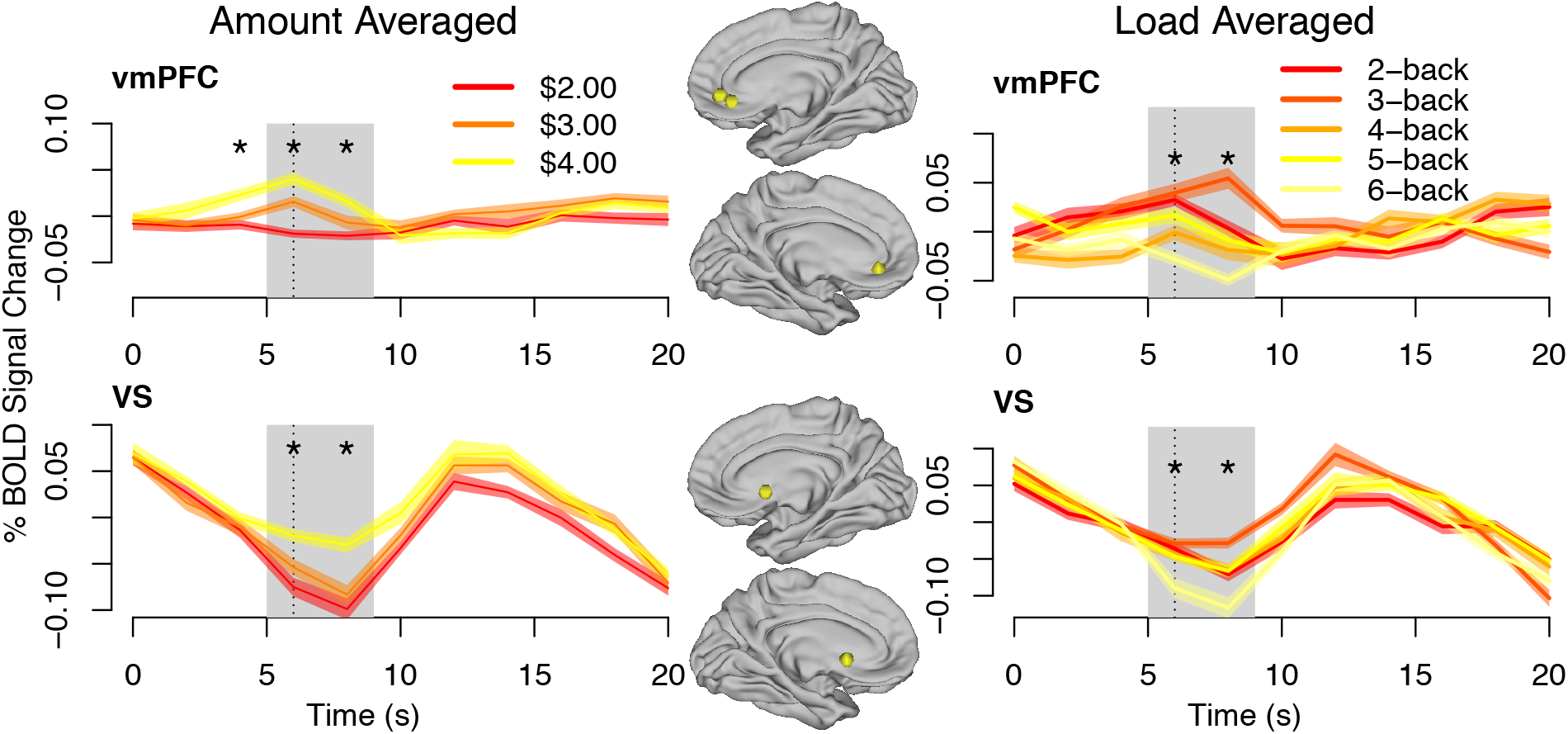
Residual timecourses in *a priori* ROIs, averaged by first offer amount or load. Error bands reflect SEM across participants. Reliability of amount and load effects in separate hierarchical multiple regression (trials and ROIs, nested within participants) at each time point indicated by * for p < 0.05. Grey region highlights 6—8 s after first offer onset; vertical dashed line indicates second offer onset.

There were some individual ROIs within the SV meta-analysis network, however, for which our data only support the encoding of single dimensions. For example, load reliably (and negatively) predicted activity in the dACC and the bilateral AI (all p’s ≤ 0.02), but reward did not (all p’s ≥ 0.18). These findings leave open the possibility that certain ROIs tracking SV here and in other studies may have reflected encoding of specific dimensions, rather than SV *per se*. There may be methodological reasons, however, why these ROIs do not track first offer amount, (e.g. low power) so particular negative results should be interpreted with caution.

### Trait Subjectivity in Value Encoding

Beyond objective dimensions like reward amount and task load, SV further implies *subjectivity* in how those dimensions are experienced. Indeed, participants varied considerably in their willingness to exert effort for reward across load and reward levels (Figure 1). To investigate subjectivity in valuation, we tested whether SV (a subjective measure) predicted valuation period activity (at 6—8 s), controlling for objective features (the amount / load ratio) in two regions that reliably tracked SV: the VS and vmPFC. In the left VS (B = 3.85×10^−2^, p = 0.03) and right (trending; B = 3.95×10^−2^, p = 0.08), SV remained a positive predictor of activity, controlling for the ratio of amount to load. The same test across all three vmPFC loci (nested within participants) was not statistically reliable (B = 1.40×10^−2^, p = 0.13), implying weaker evidence for trait subjectivity in the vmPFC.

Subjectivity may partly reflect stable, trait experience. To test for *trait subjectivity* in the experience of offers, we utilized AUC, a measure of participants’ overall tendency to accept an offer to perform the N-back for reward. Prior work has shown that COGED AUC predicts personality traits and individual differences in delay discounting, cognitive aging, and negative schizophrenia symptoms (Culbreth et al., 2016; Westbrook et al., 2013), supporting its use as a trait measure. One possible explanation for covariance across these diverse domains is reward sensitivity. Thus, we tested whether AUC predicted the effect of offer amount (reward) on SV representations.

Specifically, we fit models with amount and load jointly predicting activity in those ROIs reliably tracking SV for each participant, and then tested whether AUC predicted individual differences in fitted amount effects. Furthermore, we took advantage of the fact that we measured participants’ AUC in multiple sessions to estimate stable, trait-like tendencies to discount rewards for performing the N-back (see Supplement). In both the left (B = 2.41×10^−2^, p < 0.01) and right VS (B = 2.57×10^−2^, p = 0.06, trending), and left (B = 1.85×10^−2^, p = 0.09, trending) and right amygdala (B = 3.01×10^−2^, p < 0.01), amount effects were positively predicted by mean, cross-session AUC. Note that excluding the high amount effect participant (B_VS Amount Effect_ = 0.24; Figure 6) did not attenuate the relationship between AUC and amount effects for the remaining participants in either the left (B = 2.41×10^−2^, p < 0.01) or right VS (B = 1.35×10^−2^, p = 0.04). Furthermore, the formal interaction of AUC and amount was also a reliable predictor of BOLD signal within each of these ROIs separately, demonstrating the robustness of these effects (in a hierarchical regression model; see Supplement). This AUC-amount interaction implies that high AUC participants were more willing to perform the high-load N-back because they were more sensitive to increasing reward amounts. Indeed, the interaction in the VS and amygdala appears to be driven by higher activity for $4 offers among high versus low AUC participants (Supplement, Figure S.3). This suggests that high AUC participants perceived higher SV for $4 offers (relative to low AUC participants), via stronger encoding in the VS and amygdala, and this translated into a greater willingness to accept these high reward offers.

**Figure 6.**
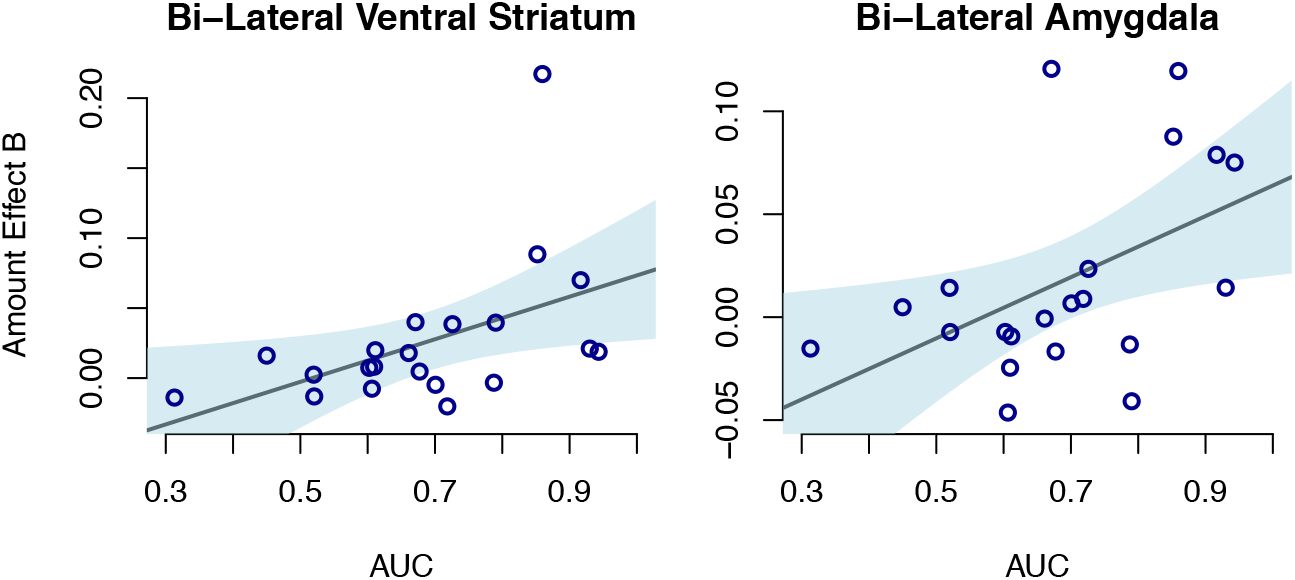
Cross-session AUC predicts the average amount effect on mean activity at 6—8 s following first offer onset in both the bilateral ventral striatum (B = 2.49×10^−2^, p = 0.03) and bi-lateral amygdala (B = 2.43×10^−2^, p = 0.02). Shaded regions show 95% CI.

### State Value Signals in the vmPFC Predict Subsequent Choice

In addition to stable, trait subjectivity, valuation may also involve state variation, from factors such as satiety or fatigue that can influence motivation (Hare et al., 2014; Rudorf and Hare, 2014). Although we did not manipulate state motivation directly, most non-catch trials were close enough to indifference for choice to be sensitive to motivational fluctuations from trial to trial. To test for such effects, we examined whether trial-wise variation in SV representations predicted choice; specifically, whether higher valuation period signal predicted high-effort choices, above and beyond the bias (*γ*) imposed by the second offer. We focused our analysis on the vmPFC, given clear SV tracking in this region, and also prior literature implicating vmPFC both in state incentive motivation, and in causally determining choice (Hare et al., 2014; 2009; 2011; Rudebeck and Murray, 2014; San-Galli et al., 2016).

During the valuation period, we found that higher vmPFC activity predicted choice of the high-load option, and that this effect was most pronounced on antibias trials (i.e., when the offer was designed to bias choice of the low-load offer). A hierarchical model with trials and 3 vmPFC ROIs nested within participants, revealed that this interaction of choice and bias significantly predicted mean vmPFC activity during the valuation period (B = 7.89×10^−2^; p < 0.01). Thus, even before participants knew what the low-load (1-back) offer amount would be, trial-wise variation in vmPFC activity predicted subsequent choice - consistent with state-dependent, causal SV representations (Figure 7). Notably, this pattern was *not* observed in either the right (B = 2.86×10^−2^; p = 0.57) or left VS (B = 5.48×10^−2^; p = 0.22), despite the VS otherwise co-varying with SV. Stronger coupling of choice to vmPFC activity is consistent with the hypothesis that while both VS and vmPFC are part of a distributed valuation network, the vmPFC serves as a final common node incorporating state motivation during decision-making (Levy and Glimcher, 2012). As shown in Figure 7, the interaction of choice and bias also reliably predicted BOLD signal later in trials with the opposite sign: activity was higher on trial in which participants selected the low-effort option. While it is possible that this reflects interesting post-decisional processes, the timing of this the interaction, pursuant to presentation of two offers and choice commitment precludes unambiguous interpretation of this effect.

**Figure 7.**
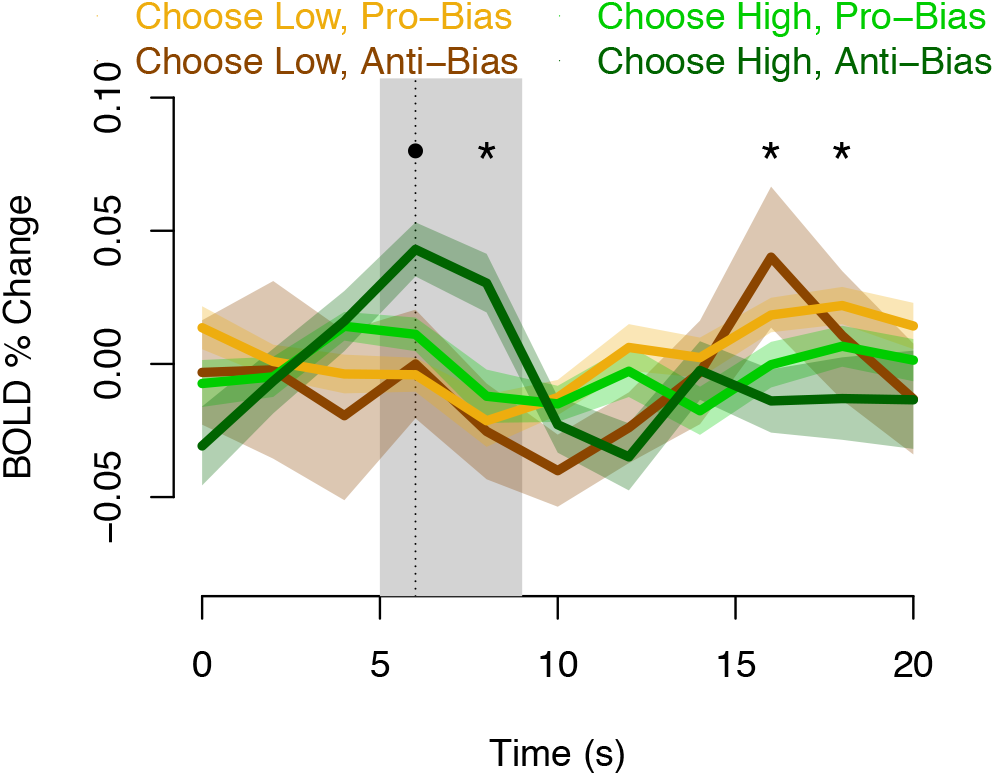
Mean residual timecourses at vmPFC loci, averaged by whether participants chose the high or low demand offer and whether the low demand offer amount biased them toward the high or low demand offer. Error bands reflect SEM across participants. Reliability of choice-bias effects in separate, fully random hierarchical multiple regressions at each time point indicated by * for p < 0.05, and • for p < 0.075. Based on a Choose Low, Pro-Bias < Choose Low, Anti-Bias < Choose High, Pro-Bias < Choose High, Anti-Bias coding scheme.

### Cognitive Demand Encoding in AI and dACC Reflects Working Memory Performance

Finally, we considered the possibility that subjective effort, and SV representations, may be related to cognitive task performance, since N-back performance correlated with discounting in the present data. Also, one recent study found that subjective effort closely tracked task performance errors (Dunn et al., 2017). Thus, we tested whether average N-back performance (the sensitivity index d′) predicted the effect of load on SV representations. We focused on the AI and dACC as these regions have been implicated previously by multiple lines of evidence, including involvement in attention and control modulation (Dosenbach et al., 2006), error awareness and processing (Klein et al., 2007), decision-making and learning about physical effort costs (Prévost et al., 2010; Skvortsova et al., 2017), and, in the case of the AI, selfreported cognitive effort ratings (Otto et al., 2014). As above, we first fit a model in which valuation period activity was jointly predicted by task load and amount separately for each participant, and then tested whether d′ predicted individual differences in load effects. Lower average N-back performance predicted more negative load effects across participants in the bilateral AI (Figure 8; B = 2.15×10^−2^; p = 0.05) though this effect was unreliable in the dACC (both p’s > 0.36). In other words, in the AI, participants with the worst average N-back performance show the strongest load effects during the valuation period (activity decreasing with increasing load). Given that this analysis collapsed performance across N-back levels, it may have been insensitive to load-specific associations in performance. Thus, we also tested the full interaction of load-specific performance and load in predicting BOLD signal 6—8s after first offer onset. In both the left AI (B = 1.22×10^−2^; p = 0.02) and right AI (B = 8.91×10^−3^; p = 0.02) and also the dACC ROI from an SV meta-analysis (Bartra et al., 2013) (B = 3.27×10^−2^; p < 0.01), and also for the dACC region identified in Study2 (Chong et al., 2017) (B = 1.46×10^−2^; p = 0.05), we found reliable, positive interactions indicating stronger load effects on BOLD signal for increasingly bad N-back performance. These interactions support the hypothesis that cognitive effort cost representations which inform the computation of SV, and thus effort discounting, are influenced by predicted task failure. That these effects obtained specifically in the AI and dACC supports their hypothesized, specialized role in processing effort costs.

**Figure 8.**
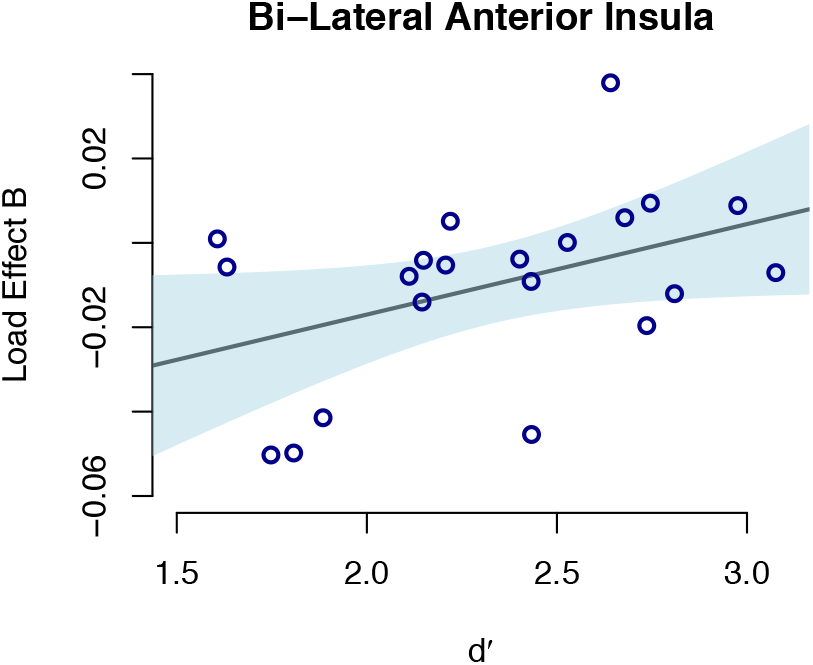
Higher mean N-back performance (measured by sensitivity index d′) predicts shallower load effects in bi-lateral AI across participants (B = 2.15×10^−2^; p = 0.05). The shaded region shows 95% CI.

### Cognitive Effort Discounting Does not Merely Reflect Task Performance

Although participants were promised payment contingent on merely repeating the N-back tasks of their choosing (not on performance), it is possible that decision-making was primarily driven by anticipated likelihood of successful performance at a given load level. In other words, an alternative interpretation of our results is rather than reflecting effort discounting and SV encoding *per se*, apparent SV encoding instead reflected concerns about performing well. This alternative account is plausible given that N-back performance covaried negatively with task load, and positively with SV. To test this alternative account, we estimated hierarchical multiple regression models to determine whether valuation period activity in the vmPFC and VS (two regions robustly encoding SV in our dataset), was predicted by SV, controlling for load level-specific N-back performance, and vice versa. The results clearly reject the alternative, performance-based interpretation. In the VS, N-back performance did not reliably predict BOLD signal variation (both p’s > 0.74), while SV did, controlling for performance (left VS: B = 3.8×10^−2^, p = 0.02; right VS: B = 4.0×10^−2^, p = 0.06). In the vmPFC, although N-back performance significantly predicted BOLD signal variation (left B = 1.4×10^−2^, p = 0.055, and right vmPFC B = 1.4×10^−2^, p = 0.046 from (Levy and Glimcher, 2012); B = 2.7×10^−2^, p = 0.02 from (Bartra et al., 2013)), SV was also a significant predictor, controlling for performance (left B = 1.8×10^−2^, p = 0.03 from (Levy and Glimcher, 2012), and right vmPFC B = 2.2×10^−2^, p = 0.06 from (Bartra et al., 2013)). These results support that while anticipated performance partially determines discounting, SV, and neural representations of SV, SV representations further reflect other state and trait factors (e.g. working memory load and reward sensitivity) determining willingness to expend cognitive effort.

## Discussion

Decisions about whether to expend cognitive effort for reward involve weighing effort costs and benefits. Indeed, subjective effort costs are reflected in demand avoidance (Kool et al., 2010; Schouppe et al., 2014b) and effort-based discounting (Botvinick et al., 2009; Chong et al., 2017; Dixon and Christoff, 2012; Massar et al., 2015; Westbrook et al., 2013). However, a central outstanding question is what mechanisms support decision-making about cognitive effort. In the current study, we used fMRI to monitor brain activity as participants made decisions about performing a high-load N-back working memory task for more money, or a low-load condition (1-back) for less money, to determine where the brain encodes the SV of rewards discounted by cognitive effort costs. Our key finding is that a putatively domain-general valuation network, centered on the vmPFC and VS, tracks cognitive effort-discounted SV. This result elucidates mechanisms of cognitive effort-based decision making, and links cognitive effort valuation closely with value-based decision-making in other domains including delay, risk, and physical effort (Bartra et al., 2013; Levy and Glimcher, 2012).

Earlier studies have shown that putative valuation regions reflect modulation of reward signals as a function of prior cognitive load, for example, or of incentive cues as a function of anticipated cognitive demands (Botvinick et al., 2009; Dobryakova et al., 2017; Nagase et al., 2018; Satterthwaite et al., 2012; Schmidt et al., 2012; Schouppe et al., 2014a; Vassena et al., 2014). These prior results imply that the domain-general valuation network should also encode SV during cognitive effort-based decision-making. Yet, to date, this has not been shown. In fact, the only two prior studies directly examining SV encoding while participants made cognitive effort decisions (Chong et al., 2017; Massar et al., 2015) largely identified regions outside the core valuation network, raising the possibility that decisions about cognitive effort rely on fundamentally different mechanisms than other cost domains. Indeed, these studies found that SV scaled with activity primarily in a dorsal, frontoparietal network typically associated with cognitive control, including the dACC / pre-SMA, dlPFC, and IPS.

Our results suggest that these two prior studies highlighted regions primarily as a function of choice difficulty, i.e., the differences in offer value during a decision trial, rather than the SV associated with the high-load offer. In particular, we identified a double dissociation, in which the putative valuation network covaried more robustly and reliably with SV than these frontoparietal regions, as a group, while conversely these frontoparietal regions were more sensitive to decision difficulty measures including offer value differences and response slowing, as a group. Note that we are agnostic as to whether frontoparietal ROIs encode Sv during offer comparison between effort-demanding alternatives. To the extent that they track differences in offer value, they may reflect differences in component value features incorporating both reward magnitude and effort demands. Indeed, the dACC / pre-SMA and ACC have been shown elsewhere to encode differences between competing alternatives in both reward and physical effort demands (Klein-Flugge et al., 2016). However, it may be that these regions are recruited *primarily* when choices are difficult, involving careful comparison between competing alternatives. This could explain why they did not reliably track first-offer SV in our study, since our design and analysis focused on SV encoding for a single offer in isolation, with first offer amount and load orthogonalized with respect to decision difficulty.

It is possible, however, to mistakenly implicate regions in encoding SV when instead they primarily encode decision difficulty, if the two are correlated. Consider, for example, general purpose control regions that are recruited more intensively to focus attention during difficult discrimination tasks. Such regions would be recruited as a function of differences in offer value, without computing SV *per se*. Thus, we reasoned, it was critical to control for decision difficulty. We designed our study to control for difficulty by ensuring that: 1) SV effects were identified during a temporally isolated valuation period in which only one offer was available, 2) offers were designed so that participants would prefer the high-demand/high-benefit option on roughly half the trials, and 3) offer attributes were fully-crossed with offer value differences (|*γ*|), guaranteeing that first-offer SV and difficulty were uncorrelated. Controlling for difficulty in this way critically revealed a double dissociation such that a core valuation network reliably tracked SV when participants evaluated a single high-load offer, but a dorsal frontoparietal network did not. Conversely, and consistent with prior studies, a dorsal frontoparietal network was recruited when value differences between offers were compared. A full account of cognitive effort-based decision-making likely involves integrated communication across multiple networks recruited for SV computation and offer comparison, particularly when choices are difficult because the offers are close in SV.

Beyond showing that a domain-general valuation network tracks first-offer SV, we further show that this network scales both positively with amount, and negatively with cognitive load. This result was critical to demonstrate that the network encoded SV rather than merely correlating with, for example, larger rewards on offer – a result which is already well-established within regions like the vmPFC and VS (Bartra et al., 2013). On an individual ROI basis, two regions – the AI and dACC – demonstrated sensitivity to cognitive load but not reward amount. This could, of course, reflect limited power to detect reward encoding in our relatively small sample. The dACC has elsewhere been shown to track both physical effort costs and reward amount, for example (Harris and Lim, 2016; Klein-Flugge et al., 2016). Also, unit recordings of monkey ACC neurons engaged in physical effort-based decision-making involves opposing signs in neighboring cells processing cost and benefit information (Kennerley et al., 2011) indicating that particular cost or benefit signals may cancel out at the resolution of fMRI. Interestingly, however, numerous lines of evidence also suggest a somewhat more specialized role for these regions in processing cognitive effort costs including their involvement in cognitive task attention and control modulation (Dosenbach et al., 2006), error awareness and processing (Klein et al., 2007), processing of affectively negative stimuli (Duerden et al., 2013), decision-making and learning about physical effort costs (Croxson et al., 2009; Kennerley et al., 2011; Kurniawan et al., 2013; Prévost et al., 2010; Skvortsova et al., 2014), self-reported cognitive effort ratings, in the case of the AI (Otto et al., 2014), and perhaps most pointedly, learning about cognitive effort costs (Botvinick, 2007; Nagase et al., 2018). Our results are thus consistent with the overarching hypothesis that the dACC and AI specialize in processing effort cost information in the service of SV computation. This interpretation is bolstered by the fact that both the dACC and AI appeared to encode cost information (load effects) as a function of individual differences in N-back task performance – two factors which we have been shown to influence cognitive effort discounting. Finally, we also note that while the AI and dACC have been shown to encode physical effort costs positively instead of negatively in several recent studies (Croxson et al., 2009; Kennerley et al., 2011; Kurniawan et al., 2013; Prévost et al., 2010; Skvortsova et al., 2014), AI and dACC activity have correlated both positively and negatively with SV across a broader value-based decision-making literature (Bartra et al., 2013) suggesting that cost sign effects (negative or positive) might be task context-dependent.

Beyond implicating a domain-general valuation network in SV encoding, our study also reveals state and trait factors which drive *subjectivity* in SV, and provides preliminary evidence about the routes by which such factors are incorporated during offer valuation. First, in the VS and amygdala, we found that shallower effort discounting (indexed with higher, cross-session AUC) predicted larger amount effects across individuals, suggesting that individuals were more willing to perform high N-back task levels in part because they were more sensitive the offered reward amounts. Trait reward sensitivity in the VS is consistent with a large literature linking VS response to reward cues with both impulsivity and other forms of psychopathology (Beck et al., 2009; Hariri et al., 2006; Plichta and Scheres, 2014). The functional coupling of the VS and the amygdala in our data is moreover consistent with close structural and functional connectivity between the regions during the processing of affective stimuli (Cardinal et al., 2002). Second, as noted above, we find evidence that SV representations reflect load-dependent performance. Specifically, in the AI and dACC, we find that negative encoding of load is stronger for those with worse N-back performance. This result is consistent with behavioral evidence that subjective effort costs are closely coupled with perceived error rates during task performance (Dunn et al., 2017). Third, we find that state (trial-by-trial) variation in SV representations in the vmPFC predict subsequent choice. Specifically, high-load offer selection was anticipated by higher-than-average vmPFC BOLD signal during the valuation period, and this was particularly true on “anti-bias” trials, in which offers biased low-load offer selection. This finding supports the hypothesis that state variation in SV, as encoded by the vmPFC, determines momentary preference (Hare et al., 2014; 2011; 2009; Rudebeck and Murray, 2014). Moreover, the pattern is also consistent with the notion that the vmPFC serves as final common pathway for economic decision-making (Levy and Glimcher, 2012), incorporating the current motivational state (Bouret and Richmond, 2010). Our fMRI result also converges with single-cell neurophysiological evidence that vmPFC activity predicts a monkey’s willingness to perform effortful instrumental tasks (San-Galli et al., 2016).

An influential hypothesis implicates the ACC/dACC in regulating cognitive control as a function of expected value, including reward benefits and effort costs (Shenhav et al., 2013). This hypothesis is motivated by prior evidence implicating the ACC in cost-benefit action selection (Croxson et al., 2009; Kennerley et al., 2009; Klein-Flugge et al., 2016; Kurniawan et al., 2010; Prévost et al., 2010; Schouppe et al., 2014a), maintaining temporally-extended action sequences (Holroyd and Yeung, 2012), and regulating cognitive control in response to cognitive demands (Botvinick et al., 2001). Our results support the hypothesis by demonstrating that information about both costs and benefits is represented in the ACC proper during offer valuation. The Expected Value of Control hypothesis does not stipulate whether the ACC/dACC would compute expected value during abstract, prospective single-offer valuation as opposed to instrumental action selection. Nevertheless, there is evidence of signed prediction-error value signaling in the ACC/dACC (reviewed in Shenhav et al., 2013), and such signaling should occur as participants are presented with unpredictable pairings of first offer amount and load. Beyond encoding cognitive costs and benefits, another prediction of the hypothesis is that the ACC/dACC region is more active when decision stakes are high, as on more difficult decision trials. Our data confirm this prediction by showing that dACC activity was higher on when participants where deciding between two offers that were close in value versus when they were far apart. In fact, the dACC was one of the few ROIs sensitive to both prospective (first offer) cognitive load and decision difficulty (the difference in first and second offer SV), a unique pattern of results which are consistent with the Expected Value of Control hypothesis.

Recent theoretical (Boureau et al., 2015; Kurzban et al., 2013; Shenhav et al., 2017) and empirical work (Dixon and Christoff, 2012; Kool and Botvinick, 2012; Westbrook et al., 2013) highlights that subjective cognitive effort costs modulate cognitive control demand avoidance, and contribute psychopathology (Cohen et al., 2001; Culbreth et al., 2016; Gold et al., 2014; Volkow et al., 2010; Westbrook et al., 2013). Using fMRI to monitor the brain while individuals considered offers to perform demanding cognitive tasks for money, we confirm that a putative, domain-general valuation network integrates cognitive costs and benefits, along with state and trait subjective factors to determine the value of cognitive effort. Importantly, and contrary to recent evidence, this network is the same one used to make decisions in other cost domains. This finding yields considerable leverage in understanding the mechanisms by which decisions about cognitive effort are made. As such, it provides a coherent set of neural targets for investigation of effort-based decision-making deficits in impaired populations, and for future development of interventions to enhance cognitive effort expenditure.

## Acknowledgements

This work was supported by grants R01 AG043461 and R21 AG058206 to T.B., and NIMH grant 1F31 MH100855−01 to A.W. Special thanks to Sarah Adams in the Cognitive Control & Psychopathology Lab at Washington University in Saint Louis for assistance with data collection.

## Author Contributions

A.W. and T.B. designed the experiment. A.W. conducted the experiments. A.W., B.L., and T.B. conducted the analyses. A.W., B.L., and T.B. wrote the manuscript.

## Methods

### CONTACT FOR REAGENT AND RESOURCE SHARING

Further information and requests for resources should be directed to and will be fulfilled by the lead contact, Andrew Westbrook.

### EXPERIMENTAL MODEL AND SUBJECT DETAILS

*Participants* Twenty-one healthy, right-handed, volunteer participants (eleven females, mean age = 21 years) recruited from the local Saint Louis community, gave informed consent as prescribed by the Institutional Review Board at Washington University. Prior to the current imaging study, all participants previously performed a separate behavioral and imaging study focused on N-back task performance, which will be the focus of a forthcoming report. Participant selection for the current study was based on ensuring both uniformly high N-back performance and sufficient individual variability in cognitive effort discounting in the included sample. Compensation for participation was provided at the rate of $25 / hour (plus additional bonus for task completion; see below).

### METHOD DETAILS

#### Apparatus and stimuli

Stimuli were presented using the Psychophysics (www.psychtoolbox.com) toolbox in MATLAB (Mathworks, Natick, MA). For each N-back load, 2 runs of 64 lower-case consonants were presented in 32 point Arial font, in one of six colors corresponding to the N-back load level: black (rgb code [0,0,0]), red [240,0,0], blue [0,0,255], purple [95,0,115], green [0,110,0], and brown [102,51,0] for the 1—6-back, respectively. Load levels were labeled by different colors, rather than by numeric load (“N”), to avoid anchoring confounds during decision-making (Chapman and Johnson, 1999). MR images were collected using a 12-channel 3-T Siemens Trio scanner (Siemens Medical Solutions USA, Inc., Malvern, PA).

#### Procedure and task design

The study was implemented in three-phases. In the first phase, following consent and screening for MR compatibility, participants performed the N-back task, completing 2 runs of each load level, in order of increasing load. In the second phase, indifference points were estimated, according to a discounting procedure detailed previously (Westbrook et al., 2013). Participants made a series of choices between repeating one of the higher N-back load level (N = 2—6) for ($2, $3, or $4) and the 1-back for amounts that varied by stepwise titration (over 5 choices per amount-load pair) until participants were indifferent between offers. In the third phase, the focus of this report, participants were scanned while performing a series of decision trials that systematically and orthogonally varied both decision difficulty and SV of the high-load offer. Over the course of 150 trials (3 amounts × 5 loads × 10 repetitions), participants made a choice between performing the high-load N-back with the given load level and reward amount, and the 1-back at a lower, variable amount. The amount offered for the 1-back was determined by the proximity parameter *γ* described in the main text. The proximity parameter *γ* was repeated twice for each value from the set {−0.4, −0.1, 0.2, 0.6} and once each from the set {−1.0, 1.0} yielding 30 catch and 120 regular trials. Each trial lasted 13 s. As noted, these decision trials began with a single offer presented in isolation for 6 s, followed by a second offer presented for up to 5.25 s, or until a decision was made. Trials were concluded with either a fixation-cross, presented after the response was made, or instead, feedback that the response deadline was missed followed by a fixation cross until the next trial. Trials in which participants did not respond in time (1.15% of trials overall) were omitted from further analyses.

Data were collected in two cohorts. In an early cohort (7 participants), T1 and T2 anatomical images were collected first, after which all decision trials were completed in a single run while 1019 functional volumes were collected. In the later cohort, to reduce discomfort, trials were broken down into 3 runs (345 volumes each); in between task runs, participants were invited to relax, motionless while T1 and T2 anatomical scans were acquired. After all images were collected, one decision trial was chosen at random, and participants were required to repeat 1 run of the chosen N-back level, after which they received the associated reward amount, as a monetary bonus added to their base compensation.

#### Imaging parameters and acquisition

T1 images were collected in 176 frames of 1×1×1 mm voxels using 2.4 s TRs, and spin-echo times of 3,080 ms, and an 8° flip angle. Anatomical T2 images were also collected in 176 frames of 1×1×1 mm voxels using 3.2 s TRs, spin-echo times of 455 ms, and a 120° flip angle. Functional imaging sequences during decision trials were collected in 4×4×4 mm voxels using a 256×256 voxel field of view, 2,000 ms TRs, 27 ms spin-echo times, and 90° flip angles.

### QUANTIFICATION AND STATISTICAL ANALYSIS

#### Imaging analyses

All DICOM images were converted to NIFTI format using the Freesurfer function mri_convert. Subsequent steps of the analysis were implemented with AFNI software functions (Cox, 1996). Brain tissue was masked using the 3dSkullstrip function, images were concatenated using 3dTcat, aligned from oblique to cardinal orientation using the 3dWarp function, and then up-sampled to 3×3×3 mm voxels and aligned across all functional runs to the first run. Parameters for registration of functional volumes with anatomical T1 images were computed for each participant separately. Precise registration was verified visually for every participant and cost functions were tailored to optimize registration for each participant. Then, parameters for warping participant-specific anatomical images to a standard MNI space (MNI152_T1_2009c+tlrc) (Fonov et al., 2011) were computed. All registration and warping parameters were concatenated using the cat_matvec function, and applied as a single transformation to aligned functional image volumes using the 3dAllineate function. Following these transformations, functional images were smoothed using an 8.0 mm FWHM kernel and the 3dmerge function.

Three distinct General Linear Models (GLMs) were fit to functional data using the 3dDeconvolve function. One GLM, designed to investigate the contrast of catch and regular trials, included stick regressors for the onset of the first offer convolved with a canonical hemodynamic response (gamma) function; this approach was used to obtain statistics in column 6 of Table 1. A second GLM, for investigating trial-wise modulation of signal by first offer SV, included finite impulse response regressors (“tent” functions in AFNI) spanning two trial epochs to capture the hemodynamic response lag (13 time points, 24 seconds), with the amplitude of each impulse response parametrically modulated by the SV of the first offer using the −stim_times_AM2 argument to 3dDeconvolve. Note that we could estimate trial-wise modulation by SV, despite fixed inter-trial intervals and overlapping hemodynamics, by exploiting the pseudo-random order and uncorrelated variation in first offer load-amount pairs. This approach was used only for Supplemental Figure S.2. A third GLM included regressors of no interest only. Residuals of this third GLM were used for most analyses, including Figures 4—8 and columns 3-5 of Table 1. All GLMs included six motion regressors, polynomial regressors suited to run duration for low-frequency drift, and a gamma-convolved stick function associated with the infrequent onset of a brief 5 s reminder menu listing all load levels in their associated color labels. In addition to motion regression, we censored all images with mean displacement ≥ 0.3 mm prior to parameter estimation to minimize the effects of high-motion transients (Siegel et al., 2013).

After fitting GLMs for individual subjects, we conducted random effects analyses. These were either on the contrast of regular and catch trials (first GLM), or on the average regression weight for the amplitude modulation 6—8 s after first offer onset (second GLM). As noted, these time points are coincident with the first 0—2 s of the second offer and are thus un-confounded by response to the second offer, accounting for hemodynamic lag. For the third GLM, we extracted and averaged residuals from all voxels lying within 6 mm radius spheres centered on loci of interest identified by prior literature. In all cases, we used exact coordinates as reported, except in the case of the amygdala, since the amygdala were slightly anterior of the center of mass reported in (Chong et al., 2017). For those loci, we used the anatomical amygdala centers as defined by (Eickhoff et al., 2005). We then conducted all statistical analyses on averaged sphere values, across trials and subjects.

For testing whether activity encodes SV across trials and subjects, we fit hierarchical linear models of residual BOLD signals (*Y*_ij_) as predicted by z-scored SV on trial *i* (*SV_ij_*), with trials nested within participants (*j*):

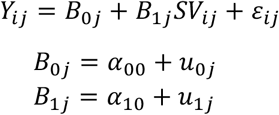

Note that for this, and all other trial-level regression analyses, we excluded trials in which participants did not respond. Also, note that for individual ROIs, we fit models in which SV predicted mean BOLD signal, after regressing motion and signals of no interest, but with no further transformation. For fitting models in which set of ROIs comprising, for example, the SV meta-analysis network, we combined all ROIs into a single outcome measure (*Y*_ij_) by first z-scoring mean residual BOLD signals at 6—8 seconds, across trials, for each ROI, then averaging these z-scores across ROIs for each participant. This method was used to give each component ROI equal weighting in quantifying the mean, network / set-level response on a given trial. For all hierarchical models, we used the *arm* package for R version 1.10-1 (https://CRAN.R-project.org/package=arm).

For testing whether sets of ROIs, as a group, predict first offer SV in a series of nested, hierarchical models, we fit the following models including either ROIs from set *A* alone (*ROIA*), then included ROIs from set *B* (*ROIB*). Note that *n_a_* refers to the total count of ROIs in set *A*, and so on.

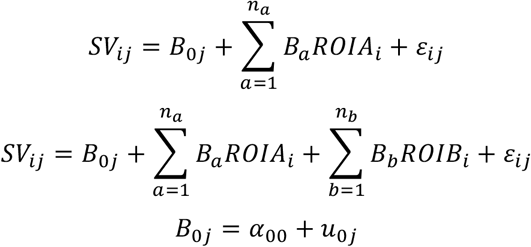

After estimating these models, we compared model fit according to *χ*^2^ distributed DIC, to ask whether the additional degrees of freedom justify the inclusion of additional ROIs as predictors in the nesting model. Note that we also conduct a parallel analysis where we swapped response slowing (inverse reaction time) for SV as the dependent variable.

Similarly, for testing whether z-scored amount and load are jointly predictive of residuals, we fit additional hierarchical linear models that replaced SV with these two terms. Note, however, that we did not model random effects for amount and load as they were fixed across participants, and also because including random effects for these predictors did not explain sufficient variance to justify the additional degrees of freedom according to a nested model comparison.

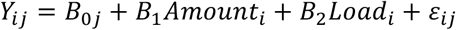

For testing whether SV predicts residuals, controlling for the z-scored ratio of amount to load (*R_A:L_*), we fit the following. Again, a nested model comparison provided evidence against estimating the random effects for the ratio predictor.

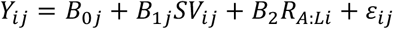

For testing whether SV predicts residuals, controlling for the ratio of amount to load in the three vmPFC ROIs (*k*), we fit the following three-level model to account for trials nested within spheres nested within participants.

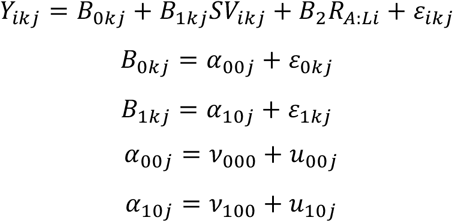

For testing whether trait AUC, at the subject level, predicts residuals as a main effect, and in interactions with amount and load, we fit the following. Note that again, amount and load are treated as fixed effects, but that AUC was a predictor of those effects at the participant level of the model.

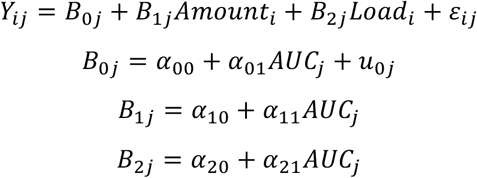

We also tested whether choice and second offer bias related to BOLD residuals present during the valuation period, fitting a model with trials, *i*, 3 vmPFC ROIs, *k*, nested within participants, *j*.

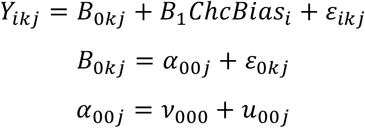

The predictor *ChcBias* was coded to incorporate two factors: 1) whether the low or high demand option was chosen; and 2) whether the amount for the low demand option was above or below indifference, thus biasing choice towards the low or high demand option, respectively. The regressor took the following values, mapping to increasing trial-wise motivation to select the high-cost, high-benefit option on a given trial. For low demand chosen, anti-bias *ChcBias* = −1.5, for low demand chosen, pro-bias *ChcBias* = −0.5, for high demand chosen, pro-bias *ChcBias* = 0.5, and for high demand chosen, anti-bias *ChcBias* = 1.5. This coding scheme was selected to emphasize the effect of higher state motivation reflected to select the high-load offer, particularly when choice was biased against this offer.

To investigate whether level-specific N-back performance (*d′_ij_*) interacted with load level in predicting BOLD signal 6—8 s after first offer onset (*Y_ij_*) on trial *i* for participant *j*, the following multi-level models were estimated in the dACC, and the bilateral AI.

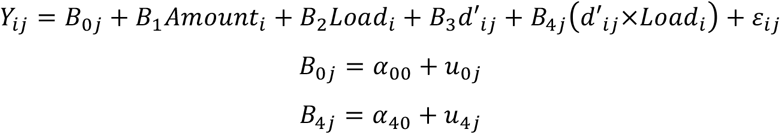

Finally, to test whether SV predicted BOLD signal 6—8 s after first offer onset, controlling for performance at that task load, we examined regression weights for the SV term in the following model. Note that random effect terms were not included for the performance predictor because they did not explain sufficient variance to justify their inclusion according to nested model comparisons.

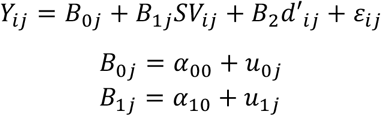

### KEY RESOURCES TABLE

| REAGENT or RESOURCE | SOURCE | IDENTIFIER |
| --- | --- | --- |
| Software and Algorithms |  |  |
| Psychtoolbox version 3.0.11 | (Brainard, 1997) | http://psychtoolbox.org/ |
| AFNI version AFNI_17.2.09 | (Cox, 1996) | https://afni.nimh.nih.gov/ |
| Freesurfer mri_convert |  | https://surfer.nmr.mgh.harvard.edu/ |
| ICBM 152 Nonlinear atlases version 2009 | (Fonov et al., 2011) | http://www.bic.mni.mcgill.ca/ServicesAtlases/ICBM152NLin2009 |
| R version version 3.4.1 | R Core Development Team (Team) | http://www.r-project.org/ |
| *arm* package version 1.10-1 | Gelman & Su (2018) | https://CRAN.R-project.org/package=arm |
| *lme4* package version 1.1-17 | Bates, Maechler, Bolker, & Walker (2015) | https://CRAN.R-project.org/package=lme4 |
| *lmerTest* package version 3.0-1 | Kuznetsova, Brockhoff, & Christensen (2017) | https://CRAN.R-project.org/package=lmerTest |
| *reshape* package version 0.8.7 | Wickham (2007) | https://CRAN.R-project.org/package=reshape |
| Functional Magnetic Resonance Imaging |  |  |
| Siemens Magnetom Trio 3T scanner | Siemens Medical Solutions, USA | http://www.siemensmriequipment.com/ |

